# ADSI-MIMO: Adaptive stain imputation with multi-input and multi-output learning for multiplex immunofluorescence imaging

**DOI:** 10.1101/2025.06.27.661891

**Authors:** Xingnan Li, Priyanka Rana, Tuba N Gide, Nurudeen A Adegoke, Yizhe Mao, Shlomo Berkovsky, Enrico Coiera, James S Wilmott, Sidong Liu

**Affiliations:** Centre for Health Informatics, Macquarie University, Sydney, NSW, Australia; Melanoma Institute Australia, The University of Sydney, Sydney, NSW, Australia; Faculty of Medicine and Health, The University of Sydney, Sydney, NSW, Australia; Charles Perkins Centre, The University of Sydney, Sydney, NSW, Australia

**Keywords:** multiplex/multispectral immunofluorescence image, generative AI, stain imputation

## Abstract

Multiplex immunofluorescence (mIF) imaging plays a crucial role in studying multiple biomarkers and their interactions within the tumour microenvironment. However, acquiring mIF images that include all desired biomarkers presents significant challenges due to the need for specialised equipment and costly reagents, which increase technical complexity, time and expense. Stain imputation offers a promising solution by synthesising target biomarker images using generative models, thereby eliminating the need for additional staining procedures. Existing deep learning-based stain imputation methods lack flexibility in generating multiple biomarker images from various input combinations. To overcome this limitation, we propose AdSI-MIMO, a novel stain imputation framework for mIF images. Our method features a multibranch deep learning architecture capable of generating multiple outputs and incorporates an adaptive progressive masking strategy to accommodate the varying combinations of input biomarkers. This approach not only improves the quality of the generated biomarker images, but also eliminates the need to train separate models for each target biomarker. We evaluated AdSI-MIMO using two datasets, including a local dataset comprising mIF images from 257 melanoma patients and a public dataset of 55 urothelial carcinoma samples. Our method achieved substantial improvements over state-of-the-art methods, particularly in the imputation of key T-cell and activation biomarkers, such as CD8 and PD-L1. Specifically, across both public and proprietary datasets, our model achieved a 18.4% improvement in Pearson Correlation Coefficient (Pearson-r) for CD8 imputation and a 48.1% improvement for PD-L1 imputation under various input biomarker configurations on the external test set.

## 1. INTRODUCTION

Multiplex immunofluorescence (mIF) imaging generates rich and multidimensional data that capture the simultaneous expression patterns of multiple biomarkers, providing valuable insights into tumour microenvironment analysis, spatial biomarker profiling, and immuno-oncology research [1]. Compared to immunohistochemistry (IHC), mIF offers several advantages by enabling a more comprehensive examination of the complex tumour microenvironment [2]. In particular, mIF allows for the detection of cell-to-cell spatial interactions, which are often associated with clinical outcomes [3, 4].

However, mIF imaging remains relatively inaccessible due to the complexities of staining and image acquisition, which requires specialised equipment and an extensive panel of biomarkers for thorough analysis [5]. Moreover, staining multiple biomarkers within a single tissue sample increases the risk of procedural failure and extends the data preparation timeline [2]. To overcome these challenges, stain imputation has emerged as a pivotal area of research. Stain imputation predicts the expression of target biomarkers and generates corresponding images by learning relationships between input and target biomarkers from reference datasets. As a result, it minimises the need for repeated staining and reduces the consumption of costly reagents.

Generative learning has emerged as a powerful approach for stain imputation, enabling accurate prediction of target biomarker expression in mIF images. A range of methods has been proposed, each supporting different input-output configurations. For instance, the Marker Imputation Model for Multiplex Images (MAXIM) [6] employs a fixed input-output structure using a conditional generative adversarial network (cGAN) [7] framework. The Stain Imputation in Multiplex Immunofluorescence Imaging (SIMIF) framework [8] introduces flexibility in input selection, while the MultimodalAttention-based virtual mIF Staining (MAS) framework [9] enables simultaneous imputation of multiple biomarkers.

Despite these advances, existing approaches still require retraining for each specific input-output configuration. This limitation underscores the need for a more adaptive and scalable framework capable of handling varying combinations of input and target biomarkers without the need for repeated model retraining.

To enhance both flexibility and effectiveness of stain imputation, we propose an adaptive framework-**AdSI-MIMO**featuring a multi-input, multi-output architecture tailored for mIF imaging. Inspired by the MultiMAE framework, which handles multiple inputs and outputs (MIMO) to leverage the partial information from multiple modalities (e.g., RGB, depth, and semantic images) [10], AdSI-MIMO adopts a similar design while introducing a novel adaptive progressive masking (APM) strategy. This strategy enables the model to accommodate varying sets of input biomarkers without requiring reconfigurations or retraining. Additionally, to accurately capture inter-biomarker dependencies, we employ a Vision Transformer (ViT) based encoder [11] for joint feature extraction. For multi-biomarker imputation, we incorporate multiple specialised decoders, each dedicated to impute a specific target biomarker. To improve image quality, we use a combination of Multi-Scale Structural Similarity Index Measure (MS-SSIM) loss [12] and *L*_1_ loss, which jointly optimise the perceptual and pixel-level fidelity of the imputed images.

In summary, our main contributions are as follows:

- MIMO architecture: The proposed framework supports flexible combinations of input and output biomarkers for mIF imaging, eliminating the need to retrain separate models for each configuration.
- APM Strategy: APM enables the model to dynamically accommodate varying input biomarker sets by selectively masking patches based on whether they are antibody-stained, significantly improving model flexibility and generalisability. Specifically, for antibiotic stains, the masking rate progressively increases throughout the training process. In contrast, for non-antibody-stained biomarker images, such as DAPI or autofluorescence (AF), the masking rate remains fixed.
- Robust imputation performance: We evaluated AdSI-MIMO on two datasets, demonstrating superior performance compared to state-of-the-art (SOTA) methods. Notably, the model maintains strong robustness and generalisability even when fewer input biomarkers are available, facilitating a more efficient and cost-effective staining process.
- Insights into biomarker panel design: Owing to its flexibility, our model enables the analysis of imputation performance across diverse input biomarker combinations. This capability offers valuable guidance for future studies in selecting optimal input biomarkers for specific targets, with the potential to reduce staining failure rates and minimise reagent costs.

## 2. RELATED WORK

### 2.1. Generative learning in histopathology imaging

With the rapid advancement of generative learning, generative models are increasingly employed to address challenges such as augmenting existing datasets to overcome data scarcity and imbalanced distribution problems [13, 14, 15]. Specifically, applications of generative models in histopathology imaging include stain normalisation and virtual staining. Stain normalisation aims to reduce variations in stain colours by mapping images to a canonical reference [16, 17], while virtual staining either simulates additional stains from a single source image [18] or generates stained images directly from unstained samples [19].

Virtual staining methods are generally categorised into supervised (paired) and unsupervised (unpaired) learning strategies. In an unsupervised setting, generative models learn to capture the style distribution of the target stain and transfer this appearance to the source images without requiring paired data [20, 21]. In contrast, supervised approaches rely on spatially registered images, typically involving unstained-stained or differently stained counterparts, allowing the model to learn a direct pixel-pixel mapping. This mapping reproduces both the colour fidelity and fine cellular morphology of the ground truth stain. Specifically, Pix2Pix-based models [22] have shown strong performance in paired virtual staining tasks [19, 23, 24]. Additionally, recent studies have demonstrated that diffusion-based models can outperform traditional GAN-based models in generating high-quality stained images [25, 26].

### 2.2. Stain imputation methods for mIF images

Stain imputation extends beyond traditional virtual staining by enabling the completion of multi-channel biomarker panels. Shaban et al. introduced MAXIM [6], a cGAN-based stain imputation method that generates the Ki67 biomarker from other available biomarkers in mIF imaging. This method incorporates a U-Net architecture [27] within the cGAN framework to model spatial relationships across biomarkers, achieving high structural similarity index measure (SSIM) scores for the imputed images. While MAXIM represent a significant advancement in stain imputation, it is constrained by a fixed input-output configuration and cannot flexibly accommodate varying biomarker combinations without full model retraining. This limitation reduces its practicality, especially given that biomarker availability and relevance often differ across experiments and tissue types. Maintaining multiple models for different configurations is also impractical for real-time use in clinical environments.

To overcome limitations of MAXIM, we proposed SIMIF [8], a more robust and flexible framework for stain imputation built on conditional Wasserstein GAN with gradient penalty (cWGAN-GP) architecture [28, 29]. This design helps mitigate common training issues such as mode collapse and vanishing gradients [13, 30], ensuring more stable training and the generation of high-quality imputed stains, even in noisy or low-signal conditions. SIMIF introduces a random channel-wise masking (RCWM) strategy that simulates scenarios in which certain biomarker inputs are absent, allowing the model to infer target stains based on the available inputs. Unlike methods that rely on fixed biomarker panels and require separate models for each input-output combination, SIMIF’s dynamic masking mechanism can accommodate diverse input configurations within a single unified framework, significantly enhancing scalability and reducing computational overhead. However, as a single-branch architecture, SIMIF is limited in its ability to generate multiple biomarkers simultaneously using a single model. Additionally, its progressive masking strategy, guided by a predefined initial probability, substantially increases training time, requiring over 1,000 epochs to impute a single biomarker from seven input channels.

To support multi-biomarker generation, MAS [9] introduces a multi-branch architecture, enabling simultaneous imputation of multiple targets. However, each input biomarker requires MAS to pre-train a separate U-Net for feature extraction, which adds architectural complexity and limits scalability-similar to the challenges seen in MAXIM. This design makes it difficult to integrate new input biomarkers, as doing so requires training new models. Moreover, by processing each biomarker independently, MAS fails to fully leverage inter-biomarker dependencies, which diminishes its imputation performance.

## 3. MATERIALS AND METHODS

### 3.1. Dataset

This study utilises two datasets, including a proprietary mIF dataset of melanoma patients [31, 32] and a publicly available dataset of urothelial carcinoma patients [33]. Both datasets include a range of biomarkers critical for predicting responses to immunotherapies [34]. Among these, CD8 and PD-L1 are of particular importance due to their essential roles in immune regulation and cancer treatment.

Quantifying CD8 and PD-L1 remains challenging because of their uneven distribution and low expression levels [35]. Therefore, we focused primarily on these two biomarkers as target biomarkers for evaluating our method, aiming to enhance the reliability and efficiency of imputing their stained images. In addition, we also assessed the model’s performance on other biomarkers to demonstrate its robustness and generalisability across a broader range of key immune biomarkers.

#### 3.1.1. Melanoma dataset

The proprietary dataset consists of 257 mIF whole slide images (WSIs), each obtained from an individual melanoma patient. Each WSI contains seven channels, corresponding to six biomarkers (CD68, SOX10, CD16, CD8, PD-L1, DAPI) and one AF channel. The images have a resolution of up to 38,000 *×* 37,000 pixels, scanned at a pixel size of 0.5 *µ*m/pixel.

These biomarkers serve different biological functions. Specifically, CD68 identifies tissue-resident macrophages and is used to assess inflammatory responses. SOX10 distinguishes parenchymal cells from immune cells. CD16 detects natural killer (NK) cells among lymphocytes. CD8 marks cytotoxic T cells. PD-L1, a key marker involved in immune regulation, is of particular interest due to its clinical relevance in immunotherapy for melanoma patients [36].

#### 3.1.2. Urothelial Carcinoma dataset

The public dataset comprises 55 mIF WSIs obtained from pre-treatment urothelial carcinoma samples. Each WSI has 15 channels, representing 13 biomarkers, with DAPI used as the base marker across three rounds of staining [33]. The images reach up to 90,000 *×* 90,000 pixels in resolution and were scanned at 0.325 *µ*m/pixel.

To ensure compatibility with our proprietary dataset, we selected 12 channels for our experiments, including core biomarkers such as CD3, CD4, CD8, CD68, SOX10, PD-L1, and DAPI. We also included additional activation markers, including LAG3, Ki67, and PD1, to further evaluate the robustness and generalisability of our model across a broader biomarker panel.

### 3.2. Data pre-processing

Images from both datasets were normalised to ensure consistent intensity levels across channels. To eliminate large background regions, tissue masks were generated using an adaptive threshold based on Otsu’s method [37]. Given the low expression levels of key biomarkers, such as PD-L1 and CD8, we combined their respective masks with the DAPI mask to guide tissue segmentation and extract the boundary coordinates of the regions of interest. This strategy effectively reduces the inclusion of empty image instances in each WSI and mitigates performance bias that may arise from imputing meaningless background regions.

The segmented tissue regions were further subdivided into 224 × 224 pixel image instances with an overlapping of 112 pixels, and each whole-slide image was capped at 2,500 image instances to keep the sample size consistent across WSIs. To enhance training efficiency, we applied a 25% sampling rate to the melanoma dataset, resulting in a final dataset of 118,923 image instances. For the urothelial carcinoma dataset, a total of 68,051 image instances were extracted.

### 3.3. Adaptive stain imputation with multi-input and multi-output (AdSI-MIMO) framework

The proposed AdSI-MIMO framework is a multi-branch generative deep learning model designed to accommodate multiple input and output biomarkers. To effectively manage varying input-output configurations, it builds upon the MultiMAE architecture and incorporates a novel APM strategy (Fig. 1). In addition, an integrated loss function combining 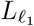 loss and MS-SSIM loss is employed to ensure preservation of both local fine-grained details and global structural integrity in the imputed images.

**Fig. 1:**
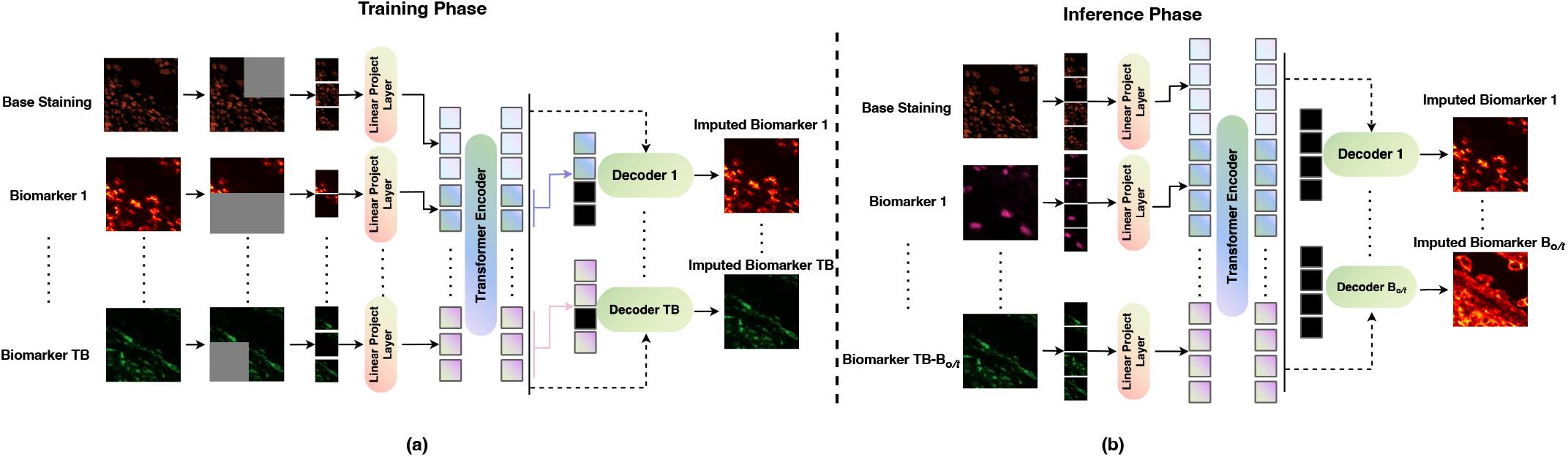
**(a)** A schematic view of the proposed AdSI-MIMO for mIF stain imputation during the training phase. **(b)** A schematic view of the proposed AdSI-MIMO for mIF stain imputation during the inference phase. Note: TB represents the number of potential target biomarkers and *B_o/t_* represents the output biomarkers.

During training, all biomarker images are provided as inputs, and the model is tasked with imputing specific target biomarkers. As illustrated in Fig. 1 (a), the biomarkers are grouped into two categories: base staining biomarkers (DAPI and AF), which are acquired through non-antibody-based fluorescence imaging and are typically easier to obtain, and antibody-stained biomarkers (e.g., CD8 and PD-L1), which serve as potential targets for imputation.

The proposed APM strategy selectively masks patches based on the biomarker category. For base biomarkers, the masking rate progressively increases using Dirichlet sampling. In contrast, antibody-stained biomarkers are masked at a fixed rate. This approach enables the model to learn how to flexibly impute target biomarkers from various subsets of available inputs.

Following patch masking, the remaining unmasked target biomarker patches are projected linearly and processed through a transformer encoder to learn a joint feature representation. These learned features serve as conditional information for each independent decoder, which then applies cross-attention to fuse the features. The self-attention block then uses these features to reconstruct the corresponding biomarker images. The number of decoders during training is equal to the number of potential target biomarkers.

In the inference phase, as illustrated in Fig. 1 (b), all base staining biomarkers are included as inputs. The encoder learns a joint feature representation from the available biomarkers, which is used to guide the decoders in reconstructing the imputed biomarker images. The total number of input biomarkers is *B* − *B_o/t_*, where *B* represents the total number of biomarkers, and *B_o/t_* is the number of target biomarkers. Each target biomarker requires a respective decoder for its reconstruction.

### 3.4. Model configuration

The proposed AdSI-MIMO framework consists of three key components: an APM block, a multi-head self-attention encoder, and a cross-self-attention (cross-self) decoder.

#### 3.4.1. Adaptive progressive masking

To accommodate varying input biomarkers, we developed an adaptive masking strategy tailored to mIF imaging. For a dataset containing *B* biomarkers, each biomarker image with a spatial dimension of *D × D* pixels, (where *D* = 224 in our study) is divided into non-overlapping patches of size *p × p*, with *p* = 16 pixels. These patches are generated through independent and trainable input adapters. Each biomarker image *b* generates (*N_pb_* = *D/p × D/p*) patches. The proposed masking strategy randomly masks patches using a fixed masking rate for non-antibody-stained biomarkers (*B_BaseStain_* (DAPI and AF)). In contrast, for potential target biomarkers, Dirichlet sampling is applied to control patch distribution between biomarkers at a progressively increasing masking rate to simulate the real-world scenarios of various signal loss in biomarker panels.

The proposed APM strategy starts with a masking probability of 30% based on the results from ablation study (*S*_*T B*0_ = *N_pB_ ×* 0.30 × *TB*) where *S*_*T B*0_ represents the number of masked patches in the dataset at epoch 0 and *TB* represents the number of potential target biomarker. This increases by 5% every 25 epochs, e.g., (*S*_*T B*25_ = *N_pB_ ×* 0.35 *× TB*), (*S*_*T B*50_ = *N_pB_*× 0.40 × *TB*), and reaches a maximum of 75% at epoch 225 (*S*_*TB*225_ = *N_pB_*× 0.75 *× TB*), which has been shown to produce optimal performance based on the MAE design [38]. Additionally, Dirichlet sampling provides masking probabilities for each potential target biomarker by generating a series of proportional values, (*d*_1_, *d*_2_ *d_T B_*), which represent the masking rate for each biomarker. These values determine the number of masked patches for each potential target biomarker (*b*_1_, *b*_2_, …*b_T B_*), computed as 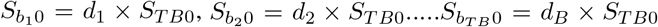 at epoch 0. For *B_BaseStain_* (DAPI and AF), we apply a fixed masking rate of 40% to prevent the encoder from over-relying on these base biomarkers when other biomarkers are heavily masked.

Specifically, all patches from each biomarker image are projected into a higher-dimensional space (768 dimensions, following the design of ViT-B [11]) as 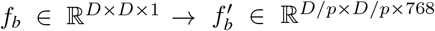, which is further reshaped to 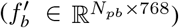. Each adapter includes a vector to encode the position information of its corresponding patch. Subsequently, following the proposed masking strategy, a subset of patches is masked within each biomarker image. The remaining unmasked patches are then concatenated across all biomarkers to form a unified input sequence, denoted as *f_Inp_* ∈ ℝ^*M*×768^, where *M* is the total number of unmasked patches. It is computed as ((*N_pb_* × *TB*) − *S_T B_*) + (*N_pb_ × B_BaseStain_* 0.60)) as illustrated in Fig. 2.

**Fig. 2:**
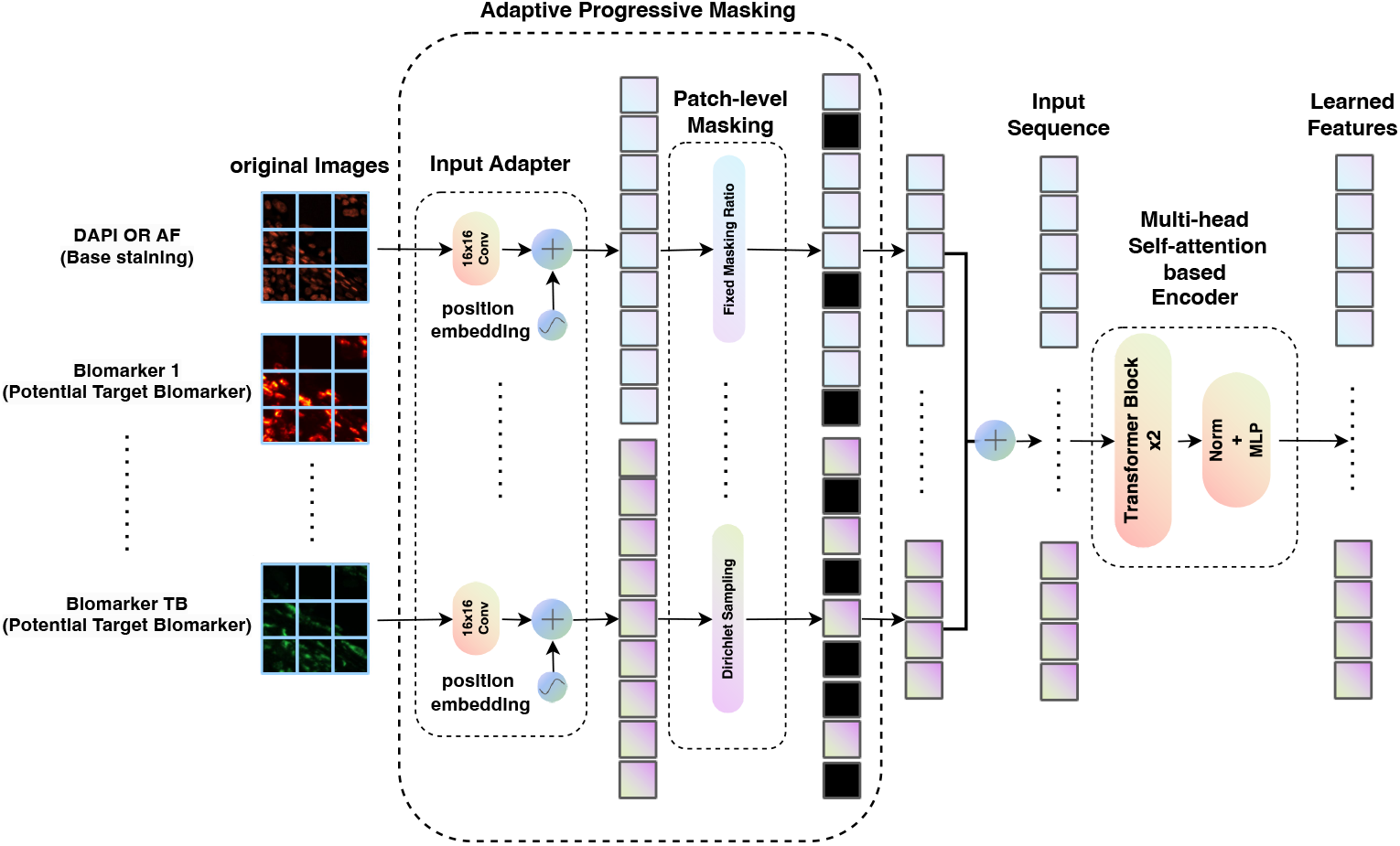
The illustration of the Adaptive Progressive Masking (APM) strategy and the Transformer based encoder in the AdSI-MIMO.

#### 3.4.2. Multi-head transformer-based encoder

We trained a ViT from scratch and used it as an encoder to extract features from the mIF images. ViT leverages the multihead self-attention mechanism of transformers, in contrast to the reliance on multiple convolutional layers, typically employed in Convolutional Neural Networks (CNNs). The encoder learns a unified feature representation of all biomarkers in the mIF image using the input sequence (*f_Inp_*) generated by the masking block, which effectively guides the subsequent decoder.

Specifically, as illustrated in Fig. 2, this unified input sequence *f_Inp_*, is fed into a multi-head self-attention module followed by the normalisation layer and a multi-layer perception (MLP) layer. In multi-head self-attention, the input is simultaneously processed across multiple attention heads, allowing the model to capture diverse relationships and patterns from different subspaces of the input representation. This process generates refined mIF image features, captures information from all biomarkers, maintains the same dimensionality, and is represented as *f_LF_* ∈ ℝw^*M* ×768^.

#### 3.4.3. Cross-self-attention decoder

In order to generate imputed biomarker images, AdSI-MIMO employs multiple independent cross-self decoders, each assigned to a target biomarker [39]. Specifically, while shared encoders would improve the model efficiency, they may compromise the ability to capture biomarker-specific features. We therefore opted to use independent decoders instead. Each decoder comprises three main modules: a cross-attention module, an MLP layer, and two self-attention modules, as illustrated in Fig. 3. Specifically, each decoder receives the biomarker image features, which constitute unmasked and masked patches 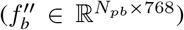 from the masking block. Moreover, it uses two additional vectors: one to encode the positional information of each corresponding patch, and another to specify the task associated with each decoder, both of dimension 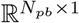.

**Fig. 3:**
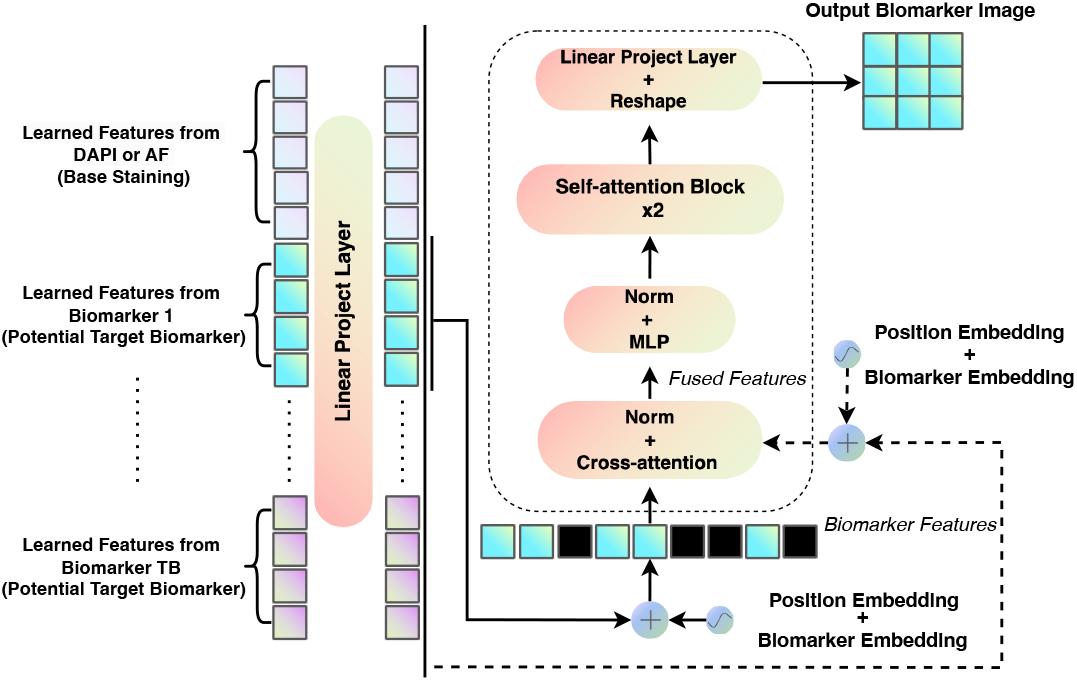
Illustration of the cross-self decoder architecture within the AdSI-MIMO framework. In the melanoma dataset, five decoders are employed, whereas in the urothelial carcinoma dataset, nine decoders are used—matching the number of potential target biomarkers.

The cross-attention mechanism [40], utilises the learned features from the encoder (*f_LF_* ∈ ℝ^*M* ×768^) as conditional information, guiding the decoder to generate target images with positional and biomarker embeddings (ℝ^*M* ×1^). This results in fused features 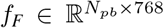. Specifically, this process is similar to self-attention, except that *Q, K*, and *V* are drawn from different sources. The *Q*, i.e., *X*_decoder_*W^Q^*, originate from the decoder, while the *K*(*X*_encoder_*W^K^*)and *V* (*X*_encoder_*W^V^*) come from the encoder. The formula for crossattention is then given by:

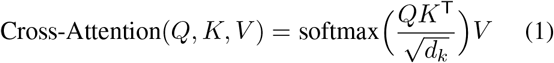

The MLP layer then adjusts the dimension of these fused features 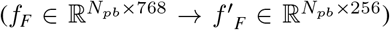 to align with the target dimensions of the imputed images. This step enables the following self-attention module to reconstruct the target biomarker images. The self-attention mechanism selectively focuses on the most informative patch embeddings and reconstructs the patches, thereby enhancing the accuracy of image reconstruction. Finally, the MLP layer maps the final features to the size of the original image (*f_output_* ∈ ℝ^*D*×D×1^). In this study, we implemented five cross-self decoders for the private dataset and nine for the public dataset.

#### 3.4.4. Loss function

To optimise model training, we integrated two loss functions, the 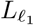 loss and the MS-SSIM loss [41]. The 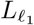 loss reduces artifacts and improves convergence, while MS-SSIM loss, as a perceptual loss, leverages characteristics of the human visual system to capture local image patterns. This combined loss function consistently improves both visual quality and the associated evaluation metrics [12]. The proposed loss function is defined as

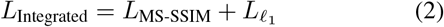

Here *L*_MS-SSIM_ loss function is defined as

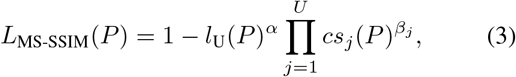

where *l_U_* (*P*) represents the luminance comparison at the coarsest scale, U captures the overall brightness consistency between the predicted and ground truth, *cs_j_*(*P*) denotes the contrast comparison at each scale *j*, evaluating the degree of alignment in local intensity variations between the two images. Additionally, *U* is the number of scales over which the structural similarity is computed, allowing the loss to account for features at multiple levels of detail. The parameters *α* and *β_j_* are weights for the luminance and contrast terms, respectively, and are set to 1 by default for simplicity. The 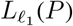 represents the loss function computed for an image instance *P*, defined as:

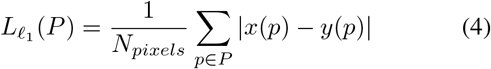

where *P* represents the image instance of pixels being evaluated, which serves as the input region for computing the loss. *N_pixels_* is the total number of pixels in the image instance, used as a normalisation factor for the loss calculation. Here, *x*(*p*) is the predicted pixel value at a specific location *p* within the image instance, and *y*(*p*) denotes the corresponding ground truth pixel value, providing a reference for comparison.

### 3.5. Evaluation metrics

Pearson Correlation Coefficient (Pearson-r) [42] measures the linear relationship between the pixel intensities of the generated image and the ground truth image as:

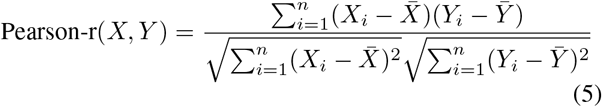

where *X_i_* and *Y_i_* represent pixel values at position *i*, while 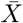 and 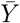 are the mean pixel values for the generated and ground truth images, respectively. The Pearson-r normalises the covariance by dividing it by the product of the standard deviations, ensuring a scale-independent measure of correlation. The result is a correlation coefficient between -1 and 1, where 1 indicates a perfect positive correlation, 0 indicates no correlation, and -1 indicates a perfect negative correlation. A high correlation value indicates a strong linear resemblance between the generated and real images. This metric is useful for assessing overall similarity in pixel intensity distributions but does not account for structural differences.

SSIM [43], on the other hand, focuses on perceived visual quality by evaluating structural information in the images, taking into account luminance, contrast, and structure. It is computed as:

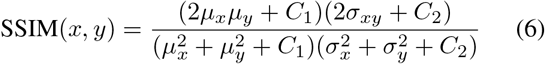

where *µ_x_* and *µ_y_* represent the mean values (luminance) of the generated and ground truth images, respectively, while 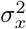 and 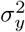 are the variances (contrast), and *σ_xy_* is the covariance (structure) between the two images. The constants *C*_1_ and *C*_2_ are used to stabilise the calculation and prevent division by zero. The formula combines these three factors, where a high SSIM score (closer to 1) indicates that the generated image has structural and perceptual qualities similar to the ground truth. This metric is widely used in image generation tasks because it aligns better with human visual perception.

Peak signal-to-noise ratio (PSNR) [44] measures the fidelity of a reconstructed image by comparing the maximum possible pixel intensity with the mean squared error (MSE) introduced by the reconstruction process. It is computed as:

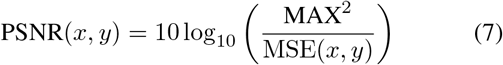

where 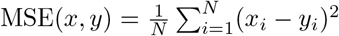 is the mean squared error between the generated image x and the ground truth image y, and MAX denotes the maximum possible pixel value in the images (e.g., 255 for 8-bit images or 1.0 for normalized images). A high PSNR value implies that the generated image has fewer distortions and is more visually similar to the ground truth, indicating a better overall reconstruction quality. This metric is widely used in image generation and restoration tasks due to its simplicity and interoperability.

### 3.6. Implementation

To prevent data leakage, we split both datasets at the WSI level. Specifically, for our private dataset, we use a stratified splitting approach, allocating 20% of the data as a test set (42 samples), 20% as a validation set (43 samples), and the remaining 60% as a training set (126 samples). We used an additional set of 46 samples, collected at a different time point, as an external test set to further evaluate the robustness of our model. In this private dataset, the inputs include all available biomarkers (CD8, PD-L1, CD68, SOX10, CD16, DAPI, and AF), while the output set excludes DAPI and AF, focusing on the remaining five biomarkers.

For the public dataset, we apply the same splitting strategy, resulting in 25 samples for training, 8 for validation, 8 for testing, and an additional 14 samples as an external test set (clinically utilising the differential treatment method). The potential output biomarkers in the public dataset include CD3, CD4, CD8, CD68, SOX10, LAG3, Ki67, PD-L1, and PD1, excluding the three DAPI channels from the iterative staining process.

All models are trained for 250 epochs with a learning rate of 1 × 10^−4^, using the Adam optimiser and a batch size of 16 on Tesla V100 GPUs with PyTorch, which takes approximately 4 days. All the configurations related to the model are fixed across these two datasets, including the APM strategy.

## 4. RESULTS

We evaluated the performance of AdSI-MIMO on the two datasets. Additionally, ablation studies were performed to assess individual contributions of each component within the proposed framework.

To further validate the model’s robustness and flexibility, we assessed its performance across varying combinations of input and output biomarkers in terms of type and number. The evaluation metrics, Pearson-r, SSIM, and PSNR, were computed at the WSI level instead of the image instance level to mitigate biases arising from the uneven image instance distribution across different WSIs. This evaluation strategy also better reflects clinical relevance by aligning with real-world practices, which prioritise slide-level interpretation over isolated image instances.

### 4.1. Comparison with state-of-the-art methods

We evaluated and compared the performance of AdSI-MIMO against current SOTA methods, including MAXIM [6], SIMIF [8], and MAS [9], for the imputation of CD8 and PD-L1 biomarkers. The results are analysed both quantitatively and qualitatively (through visual inspection).

#### 4.1.1. Quantitative analysis

We evaluated the performance of AdSI-MIMO in imputing CD8 and PD-L1 biomarkers, comparing it with existing stain imputation methods across both the melanoma (Table 1a) and urothelial carcinoma (Table 1b) datasets. For CD8, AdSI-MIMO outperformed all baseline methods in SSIM, achieving a 4.3% improvement on the melanoma dataset and a 14.6% improvement on the urothelial carcinoma dataset.

**Table 1:**
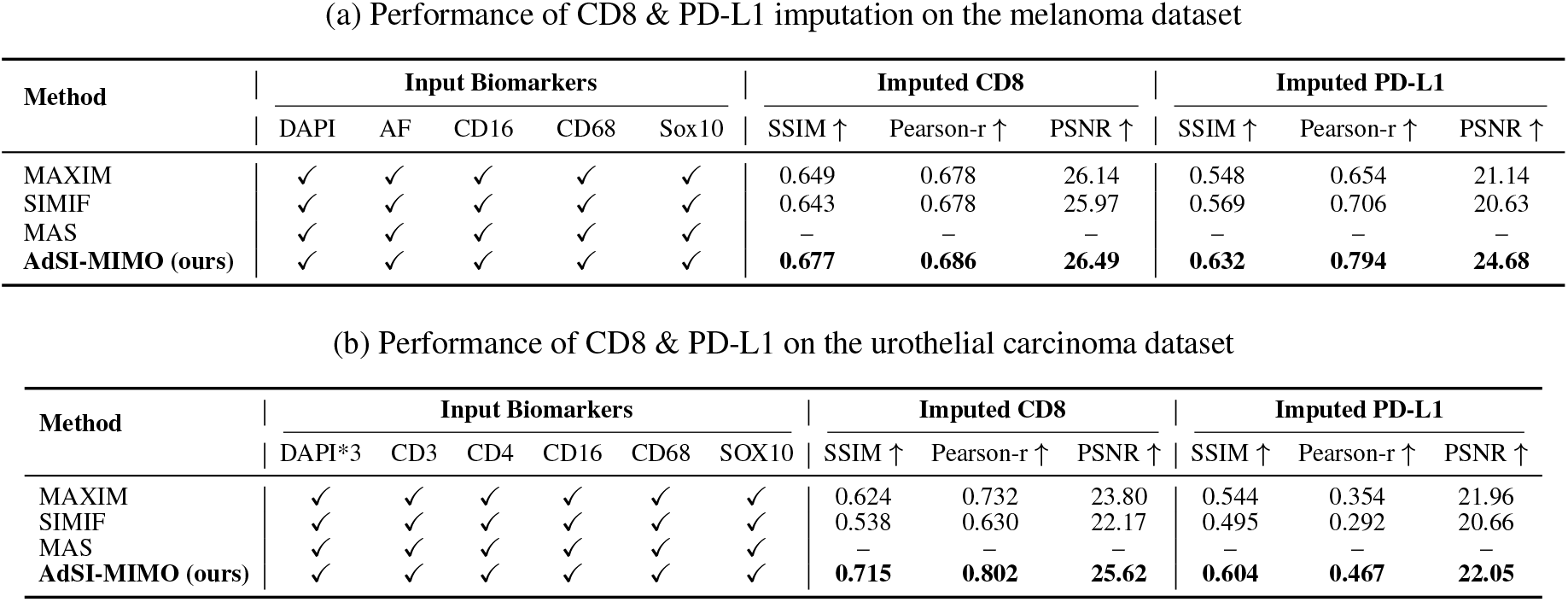
Performance comparison with all available biomarkers as input on the internal test set

(a) Performance of CD8 & PD-L1 imputation on the melanoma dataset
| Method | Input Biomarkers |  |  |  |  | Imputed CD8 |  |  | Imputed PD-L1 |  |  |
| --- | --- | --- | --- | --- | --- | --- | --- | --- | --- | --- | --- |
| | DAPI | AF | CD16 | CD68 | Sox10 | SSIM ↑ | Pearson-r ↑ | PSNR ↑ | SSIM ↑ | Pearson-r ↑ | PSNR ↑ |
| MAXIM | ✓ | ✓ | ✓ | ✓ | ✓ | 0.649 | 0.678 | 26.14 | 0.548 | 0.654 | 21.14 |
| SIMIF | ✓ | ✓ | ✓ | ✓ | ✓ | 0.643 | 0.678 | 25.97 | 0.569 | 0.706 | 20.63 |
| MAS | ✓ | ✓ | ✓ | ✓ | ✓ | — | — | — | — | — | — |
| AdSI-MIMO (ours) | ✓ | ✓ | ✓ | ✓ | ✓ | **0.677** | **0.686** | **26.49** | **0.632** | **0.794** | **24.68** |

(b) Performance of CD8 & PD-L1 on the urothelial carcinoma dataset
| Method | Input Biomarkers |  |  |  |  |  | Imputed CD8 |  |  | Imputed PD-L1 |  |  |
| --- | --- | --- | --- | --- | --- | --- | --- | --- | --- | --- | --- | --- |
| | DAPI*3 | CD3 | CD4 | CD16 | CD68 | SOX10 | SSIM ↑ | Pearson-r ↑ | PSNR ↑ | SSIM ↑ | Pearson-r ↑ | PSNR ↑ |
| MAXIM | ✓ | ✓ | ✓ | ✓ | ✓ | ✓ | 0.624 | 0.732 | 23.80 | 0.544 | 0.354 | 21.96 |
| SIMIF | ✓ | ✓ | ✓ | ✓ | ✓ | ✓ | 0.538 | 0.630 | 22.17 | 0.495 | 0.292 | 20.66 |
| MAS | ✓ | ✓ | ✓ | ✓ | ✓ | ✓ | — | — | — | — | — | — |
| AdSI-MIMO (ours) | ✓ | ✓ | ✓ | ✓ | ✓ | ✓ | **0.715** | **0.802** | **25.62** | **0.604** | **0.467** | **22.05** |

Similarly, Pearson-r showed a 1.2% increase for melanoma and a and 9.6% increase for urothelial carcinoma. For PD-L1, the proposed method achieved more substantial improvements in Pearson-r, with gains of 12.5% and 31.9% on the melanoma and urothelial carcinoma datasets, respectively.

Table 2 presents the performance of AdSI-MIMO on the external melanoma and urothelial carcinoma test sets. The results show that AdSI-MIMO consistently outperformed all baseline methods. For CD8 imputation, AdSI-MIMO achieved a 2.2% improvement in Pearson-r on the melanoma external test set and a 17.7% improvement on the urothelial carcinoma external test set. For PD-L1, the gains were even more pronounced, with improvements of 6.5% and 48.1% on the respective datasets. In addition, AdSI-MIMO demonstrated higher SSIM scores for both datasets and biomarkers, consistent with the performance trends observed on the internal test sets. Notably, the proposed model also achieved superior PSNR performance, indicating enhanced visual fidelity of the imputed biomarker images. Specifically, on the melanoma external test set, AdSI-MIMO improved PSNR by 2.5% for CD8 and 0.6% for PD-L1. On the urothelial carcinoma external test set, the PSNR improvements of 0.8% for CD8 and 0.7% for PD-L1 were observed.

**Table 2:**
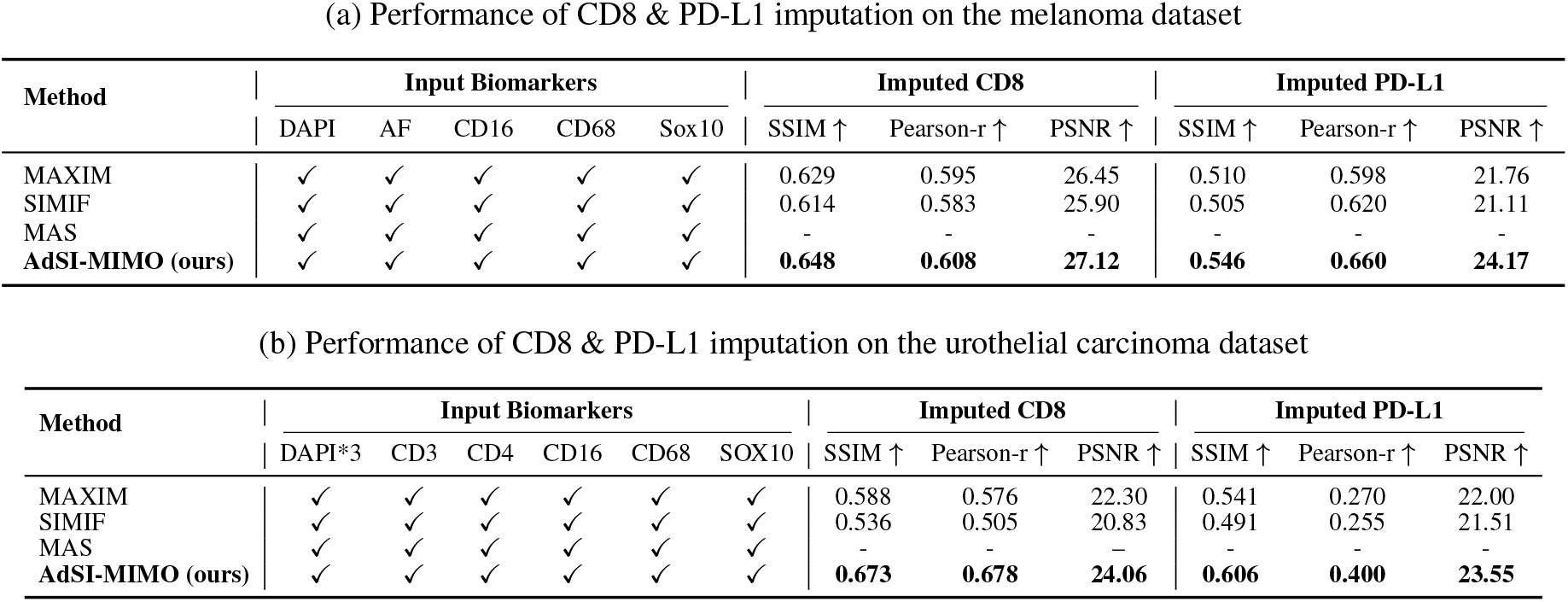
Performance comparison with all available biomarkers as input on the external test sets

We further evaluated AdSI-MIMO using only DAPI and/or AF as input, on both internal and external test sets across the two datasets, as shown in Table 3. Notably, our approach improved Pearson-r by 21.2% and 6.6% for CD8, and by 29.1% and 2.5% for PD-L1, on the melanoma (Table 3a) and urothelial carcinoma internal test sets (Table 3b), respectively. While SSIM scores for CD8 were slightly lower than those achieved by SIMIF, this can be attributed to the nature of SSIM itself, which evaluates local structural similarity using a small sliding window. SSIM is particularly sensitive to limited conditional information, especially when

**Table 3:** Performance comparison with non-antibody-stained biomarkers as input on the internal test sets

(a) Performance of CD8 & PD-L1 imputation on the melanoma dataset
| Method | Input Biomarkers |  |  |  |  | Imputed CD8 |  |  | Imputed PD-L1 |  |  |
| --- | --- | --- | --- | --- | --- | --- | --- | --- | --- | --- | --- |
|  | DAPI | AF | CD16 | CD68 | Sox10 | SSIM ↑ | Pearson-r ↑ | PSNR ↑ | SSIM ↑ | Pearson-r ↑ | PSNR ↑ |
| MAXIM | ✓ | ✓ | × | × | × | 0.393 | 0.064 | 22.24 | 0.100 | 0.011 | 16.35 |
| SIMIF | ✓ | ✓ | × | × | × | **0.545** | 0.204 | 22.37 | 0.433 | 0.430 | 19.45 |
| MAS | ✓ | ✓ | × | × | × | 0.456 | 0.274 | 22.05 | 0.392 | 0.444 | 19.17 |
| AdSI-MIMO (ours) | ✓ | ✓ | × | × | × | 0.537 | **0.332** | **23.07** | **0.486** | **0.573** | **20.80** |

(b) Performance of CD8 & PD-L1 imputation on the urothelial carcinoma dataset
| Method | Input Biomarkers |  |  |  |  |  | Imputed CD8 |  |  | Imputed PD-L1 |  |  |
| --- | --- | --- | --- | --- | --- | --- | --- | --- | --- | --- | --- | --- |
|  | DAPI*3 | CD3 | CD4 | CD16 | CD68 | SOX10 | SSIM ↑ | Pearson-r ↑ | PSNR ↑ | SSIM ↑ | Pearson-r ↑ | PSNR ↑ |
| MAXIM | ✓ | × | × | × | × | × | 0.457 | 0.207 | 20.40 | 0.312 | 0.150 | 19.37 |
| SIMIF | ✓ | × | × | × | × | × | 0.469 | 0.276 | 20.90 | 0.478 | 0.159 | 20.39 |
| MAS | ✓ | × | × | × | × | × | **0.505** | 0.456 | **21.35** | 0.464 | 0.317 | **21.20** |
| AdSI-MIMO (ours) | ✓ | × | × | × | × | × | **0.505** | **0.486** | 21.23 | **0.560** | **0.325** | 21.12 |

Pearson-r drops by more than 50% compared to scenarios with richer input data. AdSI-MIMO utilises patch-based attention mechanisms, whereas SIMIF processes the entire image holistically. Due to this difference, AdSI-MIMO tends to introduce visual discontinuities when conditional information is limited [39], leading to suboptimal SSIM scores. Despite the minor decrease in SSIM, our model achieved higher PSNR on the melanoma dataset, indicating superior overall quality compared to other SOTA models.

We also evaluated the performance of AdSI-MIMO using non-antibody-stained biomarkers as inputs on the two external test sets (Table 4). The results are consistent with patterns observed in previous experiments, further highlighting the generalisability of the proposed model for the mIF stain imputation tasks. Moreover, our model demonstrates robustness in handling the MIMO task, a capability not currently supported by any other existing model.

**Table 4:**
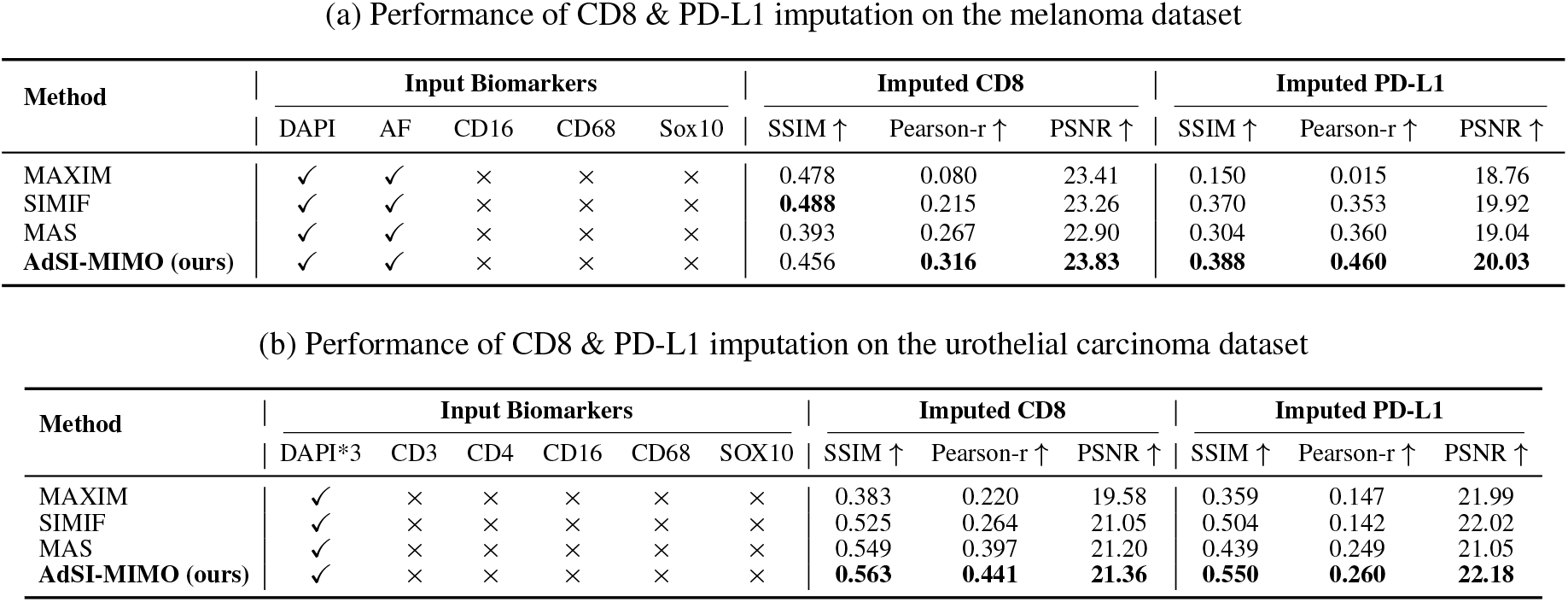
Performance comparison with non-antibody-stained biomarkers as input on the external test sets

**Table 5:**
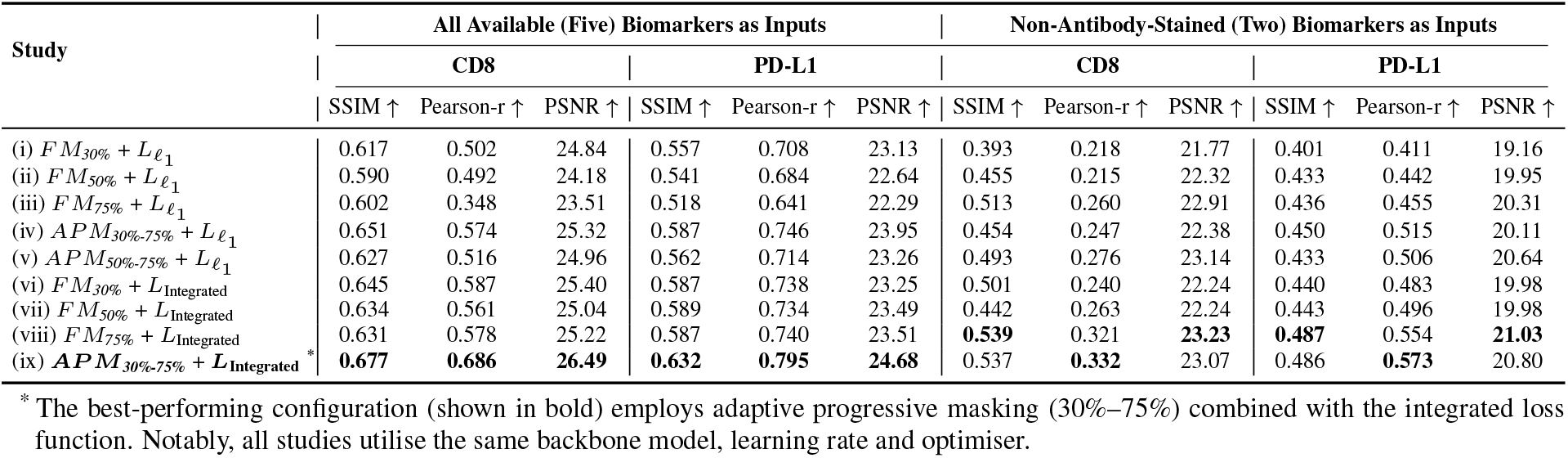
Ablation study.

#### 4.1.2. Qualitative analysis

The effectiveness of the proposed method is further demonstrated through qualitative evaluation. We visualised the imputed CD8 and PD-L1 biomarker images generated by various methods for both datasets, as shown in Fig. 4 and Fig. 5, respectively. Compared to the ground truth, AdSI-MIMO more effectively preserves fine-grained structural details-particularly in high-intensity regions-outperforming SOTA methods in visual fidelity.

**Fig. 4:**
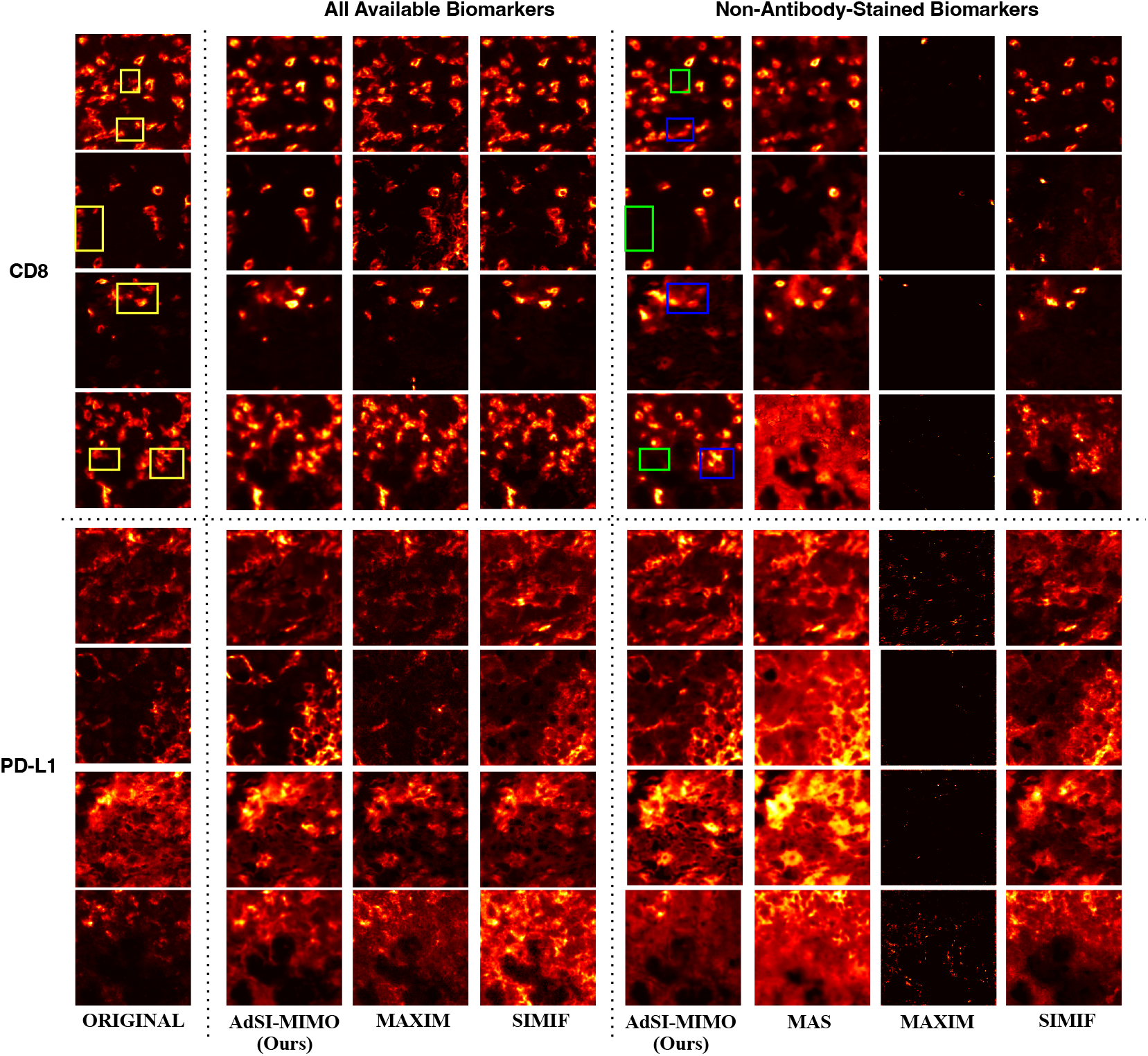
**(a)** Visualisation of original and imputed images for CD8 (top four rows) and PD-L1 (bottom four rows) in melanoma. The first two rows correspond to the internal test set, while the last two rows represent the external test set. In the AdSI-MIMO imputed images, the blue box indicates generated artifacts and the green box highlights the loss of fine details. The yellow box marks the corresponding regions in the original images.

**Fig. 5:**
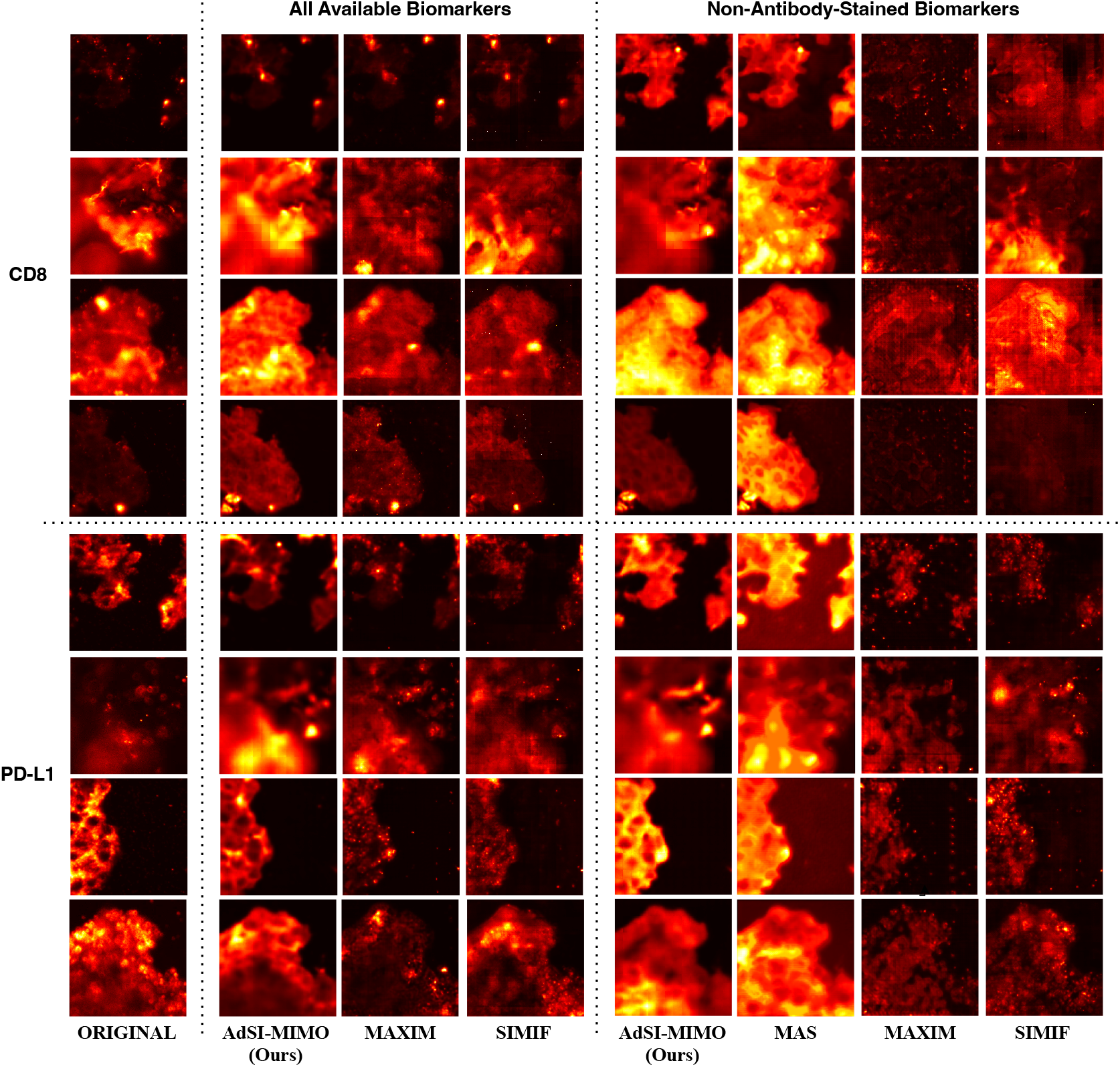
**(a)** Visualisation of original and imputed images for CD8 (top four rows) and PD-L1 (bottom four rows) in urothelial carcinoma. The first two rows correspond to the internal test set, while the last two rows represent the external test set.

For CD8 stain imputation task using all available biomarkers as input, AdSI-MIMO produces imputed images that better maintain cellular structural integrity and fluorescence distribution than baseline methods. The results appear realistic and closely aligned with the ground truth. When limited to non-antibody-stained biomarkers as input, AdSI-MIMO avoids the overly noisy or largely empty outputs observed with models such as MAXIM. However, some fine details are lost (4 green boxes), and mild artifacts (4 blue boxes) are introduced due to the restricted conditional information.

For the PD-L1 imputation, AdSI-MIMO outperform SOTA methods, particularly in reducing artifacts and preserving subtle details when all biomarkers are available. Notably, in densely clustered regions with strong fluorescence, our model more accurately captures the distribution and morphology of PD-L1 expression, resulting in visually realistic outputs. Even when only non-antibody-stained biomarkers are used, AdSI-MIMO still surpasses other methods in brightness (relative to MAS), detail retention, and overall visual quality. The imputed images contain fewer artifacts compared to CD8, indicating PD-L1 is less sensitive to the different input configurations.

Overall, AdSI-MIMO demonstrates greater stability and robustness across varying input scenarios, suggesting that model’s performance is less dependent on specific input combinations compared to existing methods.

### 4.2. Ablation studies

To evaluate the impact of the proposed APM and MS-SSIM loss function in AdSI-MIMO, we conducted a series of nine ablation studies using the melanoma dataset. Detailed results are shown in Table. 5. The compared methods are as follows:(i) fixed masking (*FM*) at 30% with the standard 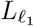 loss - maximum conditional information; (ii) *FM* at 50% with the standard loss -moderate amount of conditional information; (iii) *FM* at 75% with the standard loss - limited input (only two biomarkers); (iv) *APM* from 30% to 75% with the standard loss; (v) APM from 50% to 75% with standard loss; (vi) *FM* at 30% with the integrated loss 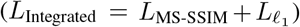; (vii) *FM* at 50% with integrated loss; (viii) *FM* at 75% with integrated loss; and (ix) APM from 30% to 75% with integrated loss. Results are based on two input configurations: all available biomarkers (five input biomarkers) and only non-antibody-stained biomarkers, with CD8 and PD-L1 as target biomarkers.

Among the fixed masking settings with standard loss (cases (i) - (iii)), the 30% masking rate excelled in the full-input setting (five biomarker images), enabling superior imputation for CD8 and PD-L1 due to the availability of sufficient conditional information. Conversely, in the limited-input setting (only two biomarkers), 75% masking yielded the best performance, as it forced the model to learn how to reconstruct the target biomarker images from minimum input-demonstrating AdSI-MIMO’s capacity to handle challenging input conditions.

Notably, the APM strategy significantly mitigated performance drop in the limited-input setting while improving results in the full-input setting. Compared to a fixed 30% masking rate, APM from 30% to 75% improved Pearson-r by 13.3% for CD8 and 25.3% for PD-L1 with two input biomarkers, and by 14.3% for CD8 and 5.4% for PD-L1 with five input biomarkers. However, CD8 performance in the limited-input setting was slightly lower than the fixed 75% masking, due to APM’s balancing of performance across both settings.

Additionally, initiating APM at 30% outperformed starting at 50%, as beginning with a higher masking rate reduced input richness in the full-input setting. This highlights the advantage of starting with a lower masking rate for more complete information, retaining the model’s ability to capture long-range dependencies.

Incorporating MS-SSIM loss (cases (vi)–(viii)) consistently enhanced the imputation quality, with notable gains in performance compared to using standard loss alone (cases (i)-(iii)). Specifically, case viii achieved the best performance for CD8 and PD-L1 imputation in terms of SSIM (0.539 and 0.487) and PSNR (23.23 and 21.03) when using two biomarkers as input. However, similar to cases (i)–(iii), it fails to maintain balanced imputation performance across different input configurations.

Among all configurations, the optimal setup for maintaining AdSI-MIMO performance across various input settings was identified as APM ranging from 30% to 75% with integrated loss (case ix). This achieved the highest overall performance in the full-input scenario: Pearson-r improvements of 16.9% for CD8 and 6.6% for PD-L1; as well as in the limited-input setting: 3.4% Pearson-r improvement for both biomarkers.

These results demonstrate the robustness of AdSI-MIMO and highlight the effectiveness of the proposed APM strategy and integrated loss function in improving imputation performance across diverse input settings.

### 4.3. Flexible input-output configurations

To demonstrate the flexibility of our model in handling diverse real-world scenarios, we tested the trained model with various input-output configurations. We summarised the corresponding performance metrics in Fig. 6 (a-c) and provided visualisation of the resulting imputed images in Fig. 6(d).

**Fig. 6:**
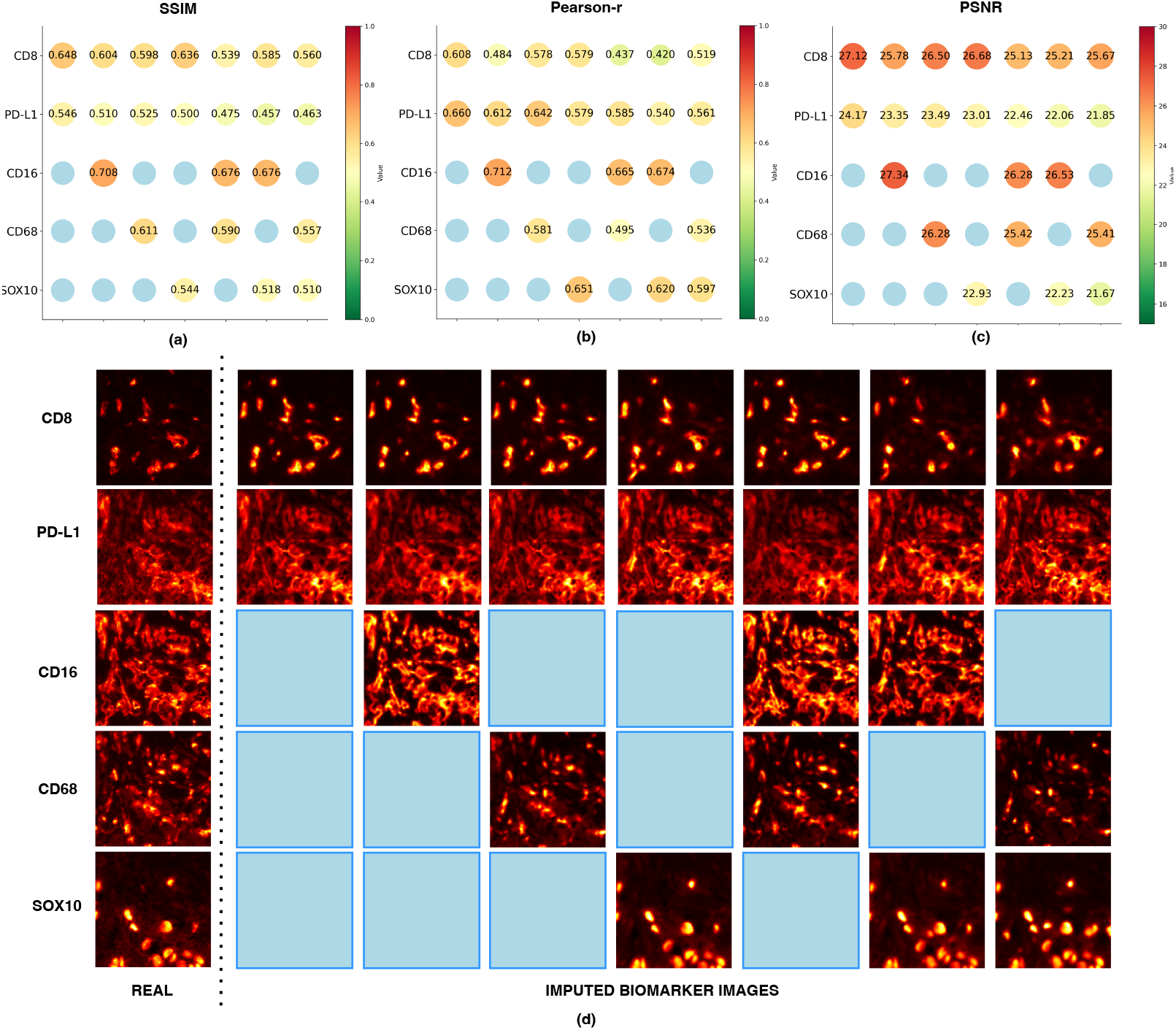
Performance of AdSI-MIMO with different input-output configurations for melanoma mIF imputation: **(a)** SSIM, **(b)** Pearson-r, **(c)** PSNR. **(d)** Visualisation of real and imputed biomarker images for different input-output combinations. Note: The x-axis represents different input configurations (blue circle or rectangle represents the corresponding biomarker used as input), while the y-axis indicates the imputation performance for the output biomarkers. Additionally, DAPI and AF, which are easily accessible biomarkers, are always included as input biomarkers by default.

In Fig. 6(a)–(c), the x-axis represents different input configurations. Specifically, DAPI and AF are always used as input biomarkers, while CD8 and PD-L1 are consistently used as target biomarkers. For CD16, CD68 and SOX10, a blue circle indicates that the biomarker was used as an input; otherwise, it was treated as an output to be imputed. The y-axis shows the imputation performance for each output biomarker. For example, in Fig. 6(a), the first column indicates that AdSI-MIMO uses CD16, CD68, SOX10, DAPI, and AF as input to impute the CD8 and PD-L1 biomarker images, achieving SSIM scores of 0.648 and 0.546, respectively.

Across all configurations, AdSI-MIMO consistently produced robust and reliable results. For example, CD16 imputation achieved SSIM ≥ 0.68, Pearson-r ≥ 0.67, and PSNR ≥ 26.3. For CD68, the model achieved SSIM ≥ 0.56, Pearson-r ≥ 0.50, and PSNR ≥ 25.4. For SOX10, the model achieved SSIM ≥ 0.51, Pearson-r ≥ 0.60, and PSNR ≥ 21.7. Overall, the summarised results demonstrated its strong performance even under varying input conditions.

We also evaluated our method to impute additional key activation biomarkers (Ki67, LAG3, and PD1) in the public dataset, demonstrating its ability to extend to a broader range of biomarkers (Fig. 7). It is observed that AdSI-MIMO can impute different activation markers while maintaining strong imputation performance and high visual quality. Specifically, across the two input settings, PD1 exhibited the greatest performance gain, while the performance of LAG3 and Ki67 remained consistent. Additionally, the Pearson-r for LAG3 reached up to 0.68, indicating that AdSI-MIMO generated highly accurate imputed images. Hence, the results are compelling as biomarkers such as CD8, PD-L1, Ki67, and PD1 provide critical information about immune infiltration and tumour aggressiveness, which can inform treatment choices, particularly in the context of immunotherapy.

**Fig. 7:**
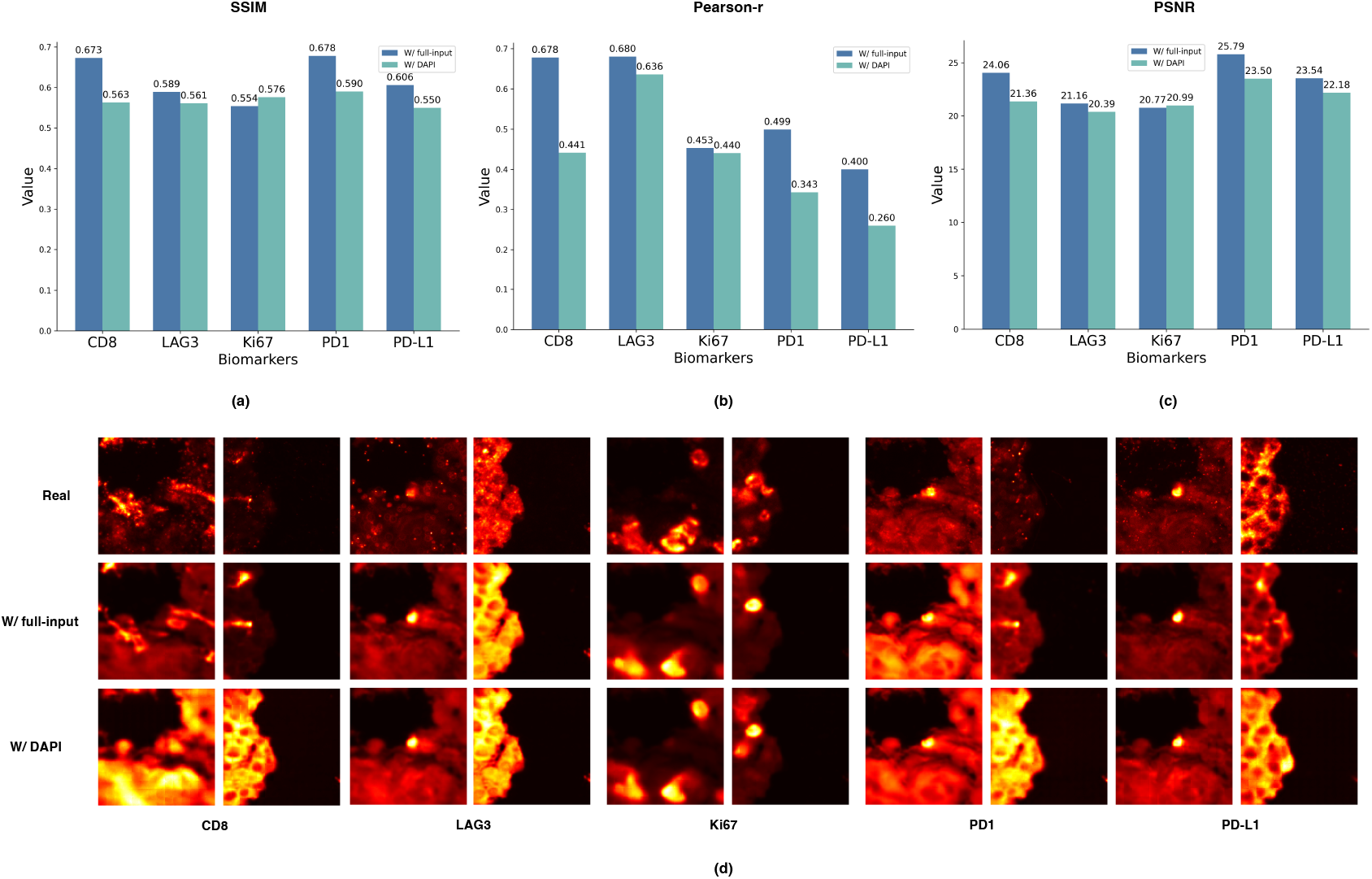
AdSI-MIMO performance with different activation markers in the urothelial carcinoma dataset, evaluated using **(a)** SSIM **(b)** Pearson-r **(c)** PSNR. **(d)** Visualisation of real and imputed activation marker images for two input-output combinations.

### 4.4. Insights into biomarker imputation performance

As illustrated in Fig. 6(a)–(c), the imputation performance decreased as the number of substituted input biomarkers increased. This observation highlights that imputation performance is highly dependent on the availability of conditional information. Notably, the performance metrics for CD8 imputation significantly declined when CD16 was excluded from the inputs for AdSI-MIMO. Specifically, Pearson-r decreased by up to 19.6% and 23.6% when using three and four biomarkers as inputs without CD16, respectively. However, this trend was not observed for the PD-L1 imputation. As depicted visually in Fig. 6(d), the quality of the imputed biomarker images aligns well with these numerical metrics. Consequently, CD16 contributes significantly to the imputation of CD8, whereas the impact of individual biomarkers is comparatively smaller for PD-L1 when certain inputs are excluded.

These findings provide valuable guidance for biomarker selection during the staining process to achieve accurate imputation according to the characteristics of the targeted biomarkers. In particular, examining the contribution of each input biomarker reveals its biomedical correlation to the imputed biomarker. As a result, histopathologists can opt to stain only the most contributive biomarkers and utilise our proposed model to accurately impute the remaining unstained targets. This strategy reduces the number of stains required, lowers reagent costs, and effectively decreases the risk of staining failures.

## 5. DISCUSSION

The primary objective of this study is to provide an adaptive, reliable generative framework, AdSI-MIMO, to impute highfidelity biomarker images from more accessible stains. We focused on imputing CD8 and PD-L1 biomarkers, given their pivotal roles in immune regulation and cancer immunotherapy, but also explored other biomarkers (Ki67, LAG3 and PD1). Comparative analyses against SOTA methods across two datasets consistently show that AdSI-MIMO outperforms existing methods. Compared to the SOTA methods, MAXIM, SIMIF and MAS, AdSI-MIMO achieves superior performance in imputing PD-L1 and CD8 biomarker images, generating outputs that more accurately reflect true staining while minimising artifacts. Its consistent performance across distinct tissue types further underscores the framework’s potential for broader adoption in diverse clinical and research oncology settings.

A key strength of AdSI-MIMO lies in its ability to learn cross-channel relationships, enabling it to effectively capture spatial and contextual dependencies among biomarkers. This results in more accurate and visually coherent imputations, particularly when the target biomarkers exhibit strong spatial correlations with available input biomarkers. This framework supports a flexible multi-input, multi-output setting, allowing the imputation of various target biomarker images from arbitrary input combinations. Our results demonstrate that imputation performance improves with an increasing number of input biomarkers. While performance naturally varies with the quantity and quality of the input, particularly when limited to non-antibody-stained biomarkers such as DAPI or AF, AdSI-MIMO remains robust and competitive even under constrained input conditions. Moreover, our proposed model significantly reduces the need for training and maintaining multiple task-specific models. This simplification is particularly valuable for pathologists managing high-throughput clinical workflows and for researchers working with largescale datasets containing diverse biomarker panels, thereby improving both scalability and operational efficiency.

Nevertheless, a few limitations remain. Despite improved performance and flexibility, the model can still produce artefacts when limited conditional information is available. This is a common issue in paired image generation. Furthermore, although cross-attention mechanisms are used to promote spatial continuity, challenges persist in generating smooth outputs under low-input settings. While results from both datasets are promising, validation on large multi-institutional cohorts is essential to assess generalisability, especially considering variability in clinical settings, such as staining protocols and scanning equipments.

Future directions include incorporating adaptive weighting or attention-based selection mechanisms to automatically prioritise the most informative biomarkers. This could inform the design of more efficient biomarker panels, reducing the costs and failure rate associated with staining while preserving high image fidelity. Furthermore, we plan to expand the model to support broader biomarkers and evaluate how imputed images affect downstream clinical tasks, such as predicting immunotherapy response. This may also involve the integration of clinical data (e.g., patient outcomes or genomic profiles) using multi-modal models, bringing stain imputation closer to clinical translation with a focus on personalised medicine.

## 6. CONCLUSION

In this study, we propose the AdSI-MIMO framework for stain imputation in mIF imaging, which features a multibranch masked autoencoder architecture, a novel Adaptive Progressive Masking (APM) strategy and an integrated loss function (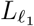 and MS-SSIM loss). It allows for flexible input-output configurations and provides a real-time virtual staining solution that is suitable for various clinical scenarios. The effectiveness of AdSI-MIMO is demonstrated by its performance in PD-L1 and CD8 imputation using two datasets. It consistently outperformed state-of-the-art methods, especially when limited to non-antibody-stained input biomarkers. In conclusion, AdSI-MIMO not only shows the ability to generate high-quality target biomarker images but also provides a rapid staining method with a flexible input-output configuration. Future work should focus on broadening its applications and testing the utility of imputed images in downstream clinical tasks.

## CRediT authorship contribution statement

Xingnan Li: Conceptualisation, Software, Methodology, Formal analysis, Investigation, Validation, Visualisation, Writing – original draft, Writing – review & editing. Priyanka Rana: Conceptualisation, Resources, Writing - review & editing, Supervision. Tuba N Gide: Data curation, Resources, Writing - review & editing, Project administration. Nurudeen A Adegoke: Data curation, Resources, Writing - review & editing. Yizhe Mao: Data curation, Resources, Writing - review & editing. Shlomo Berkovsky: Resources, Writing - review & editing. Enrico Coiera: Resources, Writing - review & editing, Funding acquisition. James S Wilmott: Data curation, Resources, Writing - review & editing, Funding acquisition. Sidong Liu: Data curation, Resources, Writing - review & editing, Supervision.

## Compliance with ethical standards

This study involves human participants and was approved by the Sydney Local Health District Human Research Ethics Committee (Protocol No. X20-0086 and 2020/ETH00426), with informed consent obtained from all participants.

## Acknowledgements

This work was supported by the NHMRC Centre of Research Excellence in Digital Health, and an NHMRC Investigator Grant (GNT2008645). Data generation was funded by the NHMRC, the Melanoma Research Alliance, Cancer Council NSW, and the CINSW Transnational Program Grant. T.N.G. is supported by a CINSW Early Career Fellowship (2020/ECF1244). J.S.W. by an NHMRC Fellowship (APP1174325), a Cancer Council NSW project grant (RG19-15), and a CINSW Translational Program Grant (TPG 2021/TPG2114).

## Data availability

Data will be made available on request.

